# H3K4me3 exhibits length-dependent deposition patterns at transcription initiation regions in *Trypanosoma cruzi* and correlates with transcriptional activity

**DOI:** 10.64898/2026.06.26.734760

**Authors:** María del Rosario López, Iván Federico Berco Gitman, Alejo Facundo Prego, Rosario Lavignolle-Heguy, Romina Trinidad Zambrano Siri, Santiago Carena, Rafael Arguello, Salomé Catalina Vílchez Larrea, Guillermo Daniel Alonso, Josefina Ocampo

## Abstract

In trypanosmatids genes, transcribed by RNA polymerase II do not have canonical promoters and are organized into directional gene clusters that mature into monocistronic transcripts by a co-transcriptional process known as trans-splicing. Even though gene expression is regulated mainly post-transcriptionally, it is currently understood that chromatin and epigenetics are also involved in this regulation. In eukaryotes, specific signals are normally required for the occurrence of an appropriate transcription initiation. Among them, trimethylation of histone H3 in lysine 4 is the most conserved signal normally detected at transcription start sites of actively transcribed genes. Unlike many model organisms, trypanosomes do not have defined promoters. Instead, transcription initiates in a bidirectional manner from dispersed regions coincident with divergent strand switch regions located between directional gene clusters **(DGCs)**. In *T. cruzi*, H3K4me3 was observed at the origins of transcription coincident with divergent strand switch regions **(dSSRs)** in epimastigotes, but it has not been mapped throughout the whole genome at base-pair resolution or in other life stages so far.

Here, we set up the CUT&RUN technique for *T. cruzi* epimastigotes and trypomastigotes. Consistent with a predominant post-transcriptional regulation along the life cycle, we did not find significant differences between life stages. We corroborated that H3K4me3 is enriched at dSSR adjacent to actively expressed DGCs. Moreover, we noticed that this histone mark exhibits different patterns that correlate with the genomic span of the transcription initiation regions and with transcriptional activity. Furthermore, we unveiled that the most actively transcribed DGCs are associated with shorter dSSRs and are located within the core compartment of the genome displaying a more accessible chromatin.

## Background

The detection of post-translational modifications **(PTMs)** of histones and its interacting networks founded the bases of the histone code (Strahl and Allis, 2000). This set of rules implies that the occurring histone PTMs determine changes in chromatin organization that can affect gene expression, leading to different phenotypical outcomes.

*Trypanosoma cruzi’s* genome is composed of a core compartment and a disruptive compartment that differentiate from each other by harboring different kinds of genes having different GC content (Berná et al., 2018). While the core compartment is rich in conserved genes, in the disruptive compartment, genes involved in virulence and immune system evasion prevail.

These parasites have a complex life cycle, alternating between an insect vector, and a mammalian host. Throughout this cycle, the parasite needs to adapt to the environmental fluctuations and suffers several morphologic changes that require modulation of gene expression. Within the insect vector, *T. cruzi* is present as epimastigotes, which possesses replication capacity. In the distant portion of the intestine of the insect vector, it differentiates to metacyclic trypomastigotes, which are the infective forms and are characterized by the absence of cell division. Within the host cells the parasite resides as amastigotes, replicative forms which must change into trypomastigotes to re-start the cycle (Tyler and Engman, 2001). Another peculiarity is that genes are organized into directional gene clusters **(DGC)**. These DGC are transcribed by RNA polymerase II (Pol II) into polycistronic units that mature into monocistronic transcripts by a co-transcriptional process known as trans-splicing (Matthews et al., 1994). Despite gene expression being regulated mainly post-transcriptionally, it is currently understood that chromatin and epigenetics are also involved in this regulation. With the advent of more sensitive technologies employed in proteomic and genomic studies, and with the improvement of genome assemblies upon the application of long read technologies, several epigenetic marks have been unveiled (Ocampo et al., 2025). However, despite unmasking the role of some epigenetic marks, most of them are still unexplored.

In eukaryotes, specific signals are normally required for the occurrence of an appropriate transcription initiation. These cues are provided by DNA sequence or epigenetic marks such as histone PTMs or histone variants. Unlike many model organisms, trypanosomes do not have defined promoters. Instead, transcription initiates in a bidirectional manner from dispersed regions coincident with divergent strand switch regions **(dSSRs)**, located between DGCs and terminates at convergent strand switch regions **(cSSRs)**. More recently, the presence of a short sequence with a promoter role was reported to be required for transcription initiation of protein-coding genes (Carvalho de Lima et al., 2024; Cordon-Obras et al., 2022; Martínez-Calvillo et al., 2003; Respuela et al., 2008; Wedel et al., 2017). Resembling the epigenetic marks found at the transcription start sites in model eukaryotes, the presence of H2A.Z variant and some histone PTMs at dSSRs were also described in trypanosomes (McDonald et al., 2022; Respuela et al., 2008; Siegel et al., 2009).

Trimethylation of histone H3 in lysine 4 **(H3K4me3)** is the most conserved histone PTM among eukaryotes and it is also the most studied one. It is normally detected at transcription start sites of actively transcribed genes and its signal intensity correlates with transcription levels (Howe et al., 2017). Moreover, the levels and patterns of H3K4 methylation are highly regulated with implications in development and cell viability (Wang and Helin, 2025). Even though trypanosomatids histones are very divergent in sequence and some histone PTMs are specific of this lineage, the N-terminal tail of histone H3 has some conserved residues and PTMs, including H3K4me3 (Ocampo et al., 2025). Genome-wide studies have shown that H3K4me3 is enriched at dSSR in *T. brucei* and *Leishmania* (Diotallevi et al., 2025; McDonald et al., 2022; Wright et al., 2010). In *T. cruzi*, a similar result was observed for a subset of chromosomic regions in epimastigotes, but it has not been mapped throughout the whole genome or in other life stages so far (Respuela et al., 2008).

In this work, we set up the CUT&RUN technique for *T. cruzi* epimastigotes and trypomastigotes. We corroborated that H3K4me3 is mainly enriched at the dSSRs with variations in intensity and patterns along the genome. Consistent with a predominant post-transcriptional regulation along the life cycle, we did not find significant differences between epimastigotes and trypomastigotes. Additionally, when H3K4me3 peaks are contrasted with RNA-seq signals, the strongest ones are next to highly expressed DGCs, suggesting that this epigenetic mark is involved in a regulatory mechanism. Remarkably, we found that H3K4me3 displays different patterns, and they are related to the extension of the dSSRs and to the transcriptional activity. Finally, we observed that the strongest and more defined H3K4me3 peaks belong to the core compartment of the genome coincident with a more accessible chromatin.

## Results

### H3K4me3 is enriched at dSSRs in *T. cruzi* epimastigotes and trypomastigotes

Although gene expression in trypanosomes is mainly regulated post-transcriptionally, it is well understood that chromatin and epigenetics fine-tune this process (Saha, 2020). In these organisms, there is a plethora of histone PTMs that are unique to this parasite. However, H3K4me3 is an almost universal epigenetic mark associated with transcription initiation (Ocampo et al., 2025). With the aim to better understand how transcription by Pol II initiates in the polycistronic units of *T. cruzi*, we pursued the detection of H3K4me3 throughout the genome of Dm28c strain. Using a specific anti-H3K4me3 antibody and a non-specific IgG as a control we adapted for the first time the CUT&RUN technique (Skene and Henikoff, 2017) for epimastigotes, in exponentially growth phase, and trypomastigotes.

We aligned the paired-end reads obtained for duplicated CUT&RUN experiments for epimastigotes and trypomastigotes to the recently published T2T version of Dm28c genome (Greif et al., 2025).

In trypanosomes, genes are organized in DGCs which are transcribed by Pol II from dSSRs to cSSRs (Fig. 1A). Detailed inspection of the peaks called with MACS2 unveiled that most of the H3K4me3 signals are located inside DGCs, but the most prominent ones localize at dSSRs. Besides, there is a small number of weak peaks detected at or nearby cSSRs, and some strong ones in other regions of the genome whose significance remains to be explored (Supplemental Fig. S1). This trend is observed both in epimastigotes and trypomastigotes. A full list of peaks detected at those locations is provided in Supplemental Table S1.

**Figure 1.**
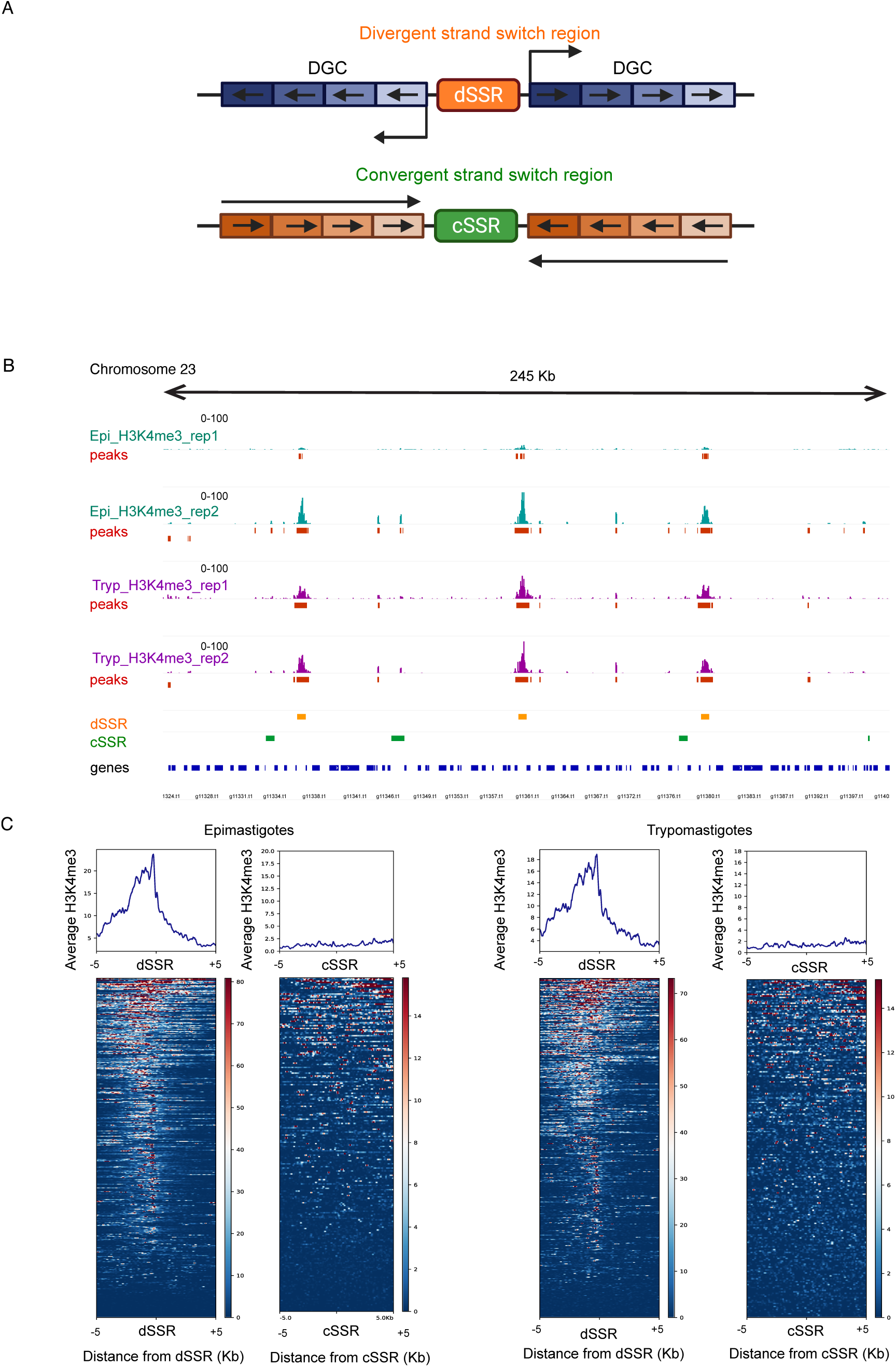
H3K4me3 is prominent at dSSRs in epimastigotes and trypomastigotes. **A)** Schematic representation of dSSRs and cSSRs demarking the directional gene clusters (DGCs). **B)** Genomic snapshot for a region of chromosome 23 of the T2T Dm28c genome displaying H3K4me3 tracks for replicated experiments in epimastigotes (cyan) and trypomastigotes (purple) with their respective narrow peaks (red) generated with MACS2. Annotated coordinates for dSSRs (orange) and cSSRs (green) for the used genome are included as a reference. **C)** Average H3K4me3 density (top panels) and heatmaps (bottom panels) relative to the boundaries of the dSSRs (left) or cSSRs (right) in a 5 Kb window for a representative replicate of epimastigotes and trypomastigotes. In the heatmaps genes are sorted according to the H3K4me3 signal in a descending order. For those represented relative to cSSRs, annotations for DGCs shorter than10Kb were removed to avoid the misleading observation of the neighbor dSSRs. Red: high signal, Blue: low signal, Green: missing data.

Additionally, genomic inspection with integrative genome viewer (IGV) allowed us to observe that the most pronounced peaks called for H3K4me3 are reproducible between replicates. In addition, consistent with previous reports, we corroborated that the strongest H3K4me3 marks were located at dSSRs and that weaker ones appeared inside the DGCs. (Fig. 1B). These observations are a common feature for the parasite stages under study.

To analyze global trends, we mapped H3K4me3 signals relative to the 5′ and 3′ boundaries of both dSSRs and cSSRs. For simplicity, these categories are used as reference points, encompassing both edges of their respective sequences. Average representation of H3K4me3 signals and heatmaps relative to dSSRs showed that H3K4me3 is enriched at the reference point in most of the genome for both life forms of the parasite. Conversely, no enrichment is observed at cSSRs, neither in epimastigotes nor in trypomastigotes (Fig. 1C and Supplemental Fig. S2A). We observed that a noticeable H3K4me3 signal is detected at most dSSRs along the genome; however, the intensity is not even among them, suggesting that transcription might not be equal for every DGC. For representations relative to cSSRs, we removed those annotated between DGCs shorter than 10 Kb to avoid the misleading observation of the neighbor dSSRs. An example of the occurrence of cSSR and dSSR within short proximity is illustrated in Supplemental Fig. S2B. Consistent with the average representation, no substantial H3K4me3 enrichment is observed at any cSSRs throughout the heatmaps.

Furthermore, to assess the conservation of H3K4me3 distribution between epimastigotes and trypomastigotes, we compared its signal across dSSRs in both life stages, using the average of two biological replicates, and represented the result into a scatterplot. Consistent with the predominant post-transcriptional regulation of gene expression, H3K4me3 peak distribution showed a high degree of similarity between life stages, with a Spearman correlation coefficient of 0.72 (Supplemental Fig. S2C). A full list containing the called peaks for replicated experiments in both life stages is provided in Supplemental Table S2.

Overall, we confirmed previous observations that H3K4me3 peaks are mainly located at dSSRs in *T. cruzi* epimastigotes, and we clarified that peaks at dSSRs were the most prominent ones, but it was not an exclusive location. Moreover, we report a conserved layout of H3K4me3 throughout the genome between epimastigotes and trypomastigotes. These observations suggest that, despite lacking canonical promoters, transcription initiation in trypanosomes is accompanied by epigenetic signposts which are preserved in the analyzed life stages of the parasite.

### H3K4me3 is prominent at highly transcribed genes and exhibits different patterns that correlate with the extension of the dSSRs

In model eukaryotes, H3K4me3 is often associated with active transcription, either by being highly enriched at TSSs or at active enhancers (Guenther et al., 2007; Shen et al., 2016). In *T. cruzi*, dSSRs work as transcription initiation regions (Carvalho de Lima et al., 2024; Respuela et al., 2008). To evaluate if this association is also true for these trypanosomes, we monitored mRNA levels using stranded RNA-seq in epimastigotes and trypomastigotes and compared the output signal with the H3K4me3 peaks. By exploring different genomic regions on IGV, we observed that the most prominent peaks of H3K4me3 are normally adjacent to, but not overlapping, mRNA signals suggesting a regulatory role for this epigenetic mark in both life stages of the parasite (Supplemental Fig. S3A). Additionally, individual inspection of H3K4me3 profiles for dSSRs with different extensions revealed that short dSSRs exhibit a defined peak as shown in S3A top panel, whereas longer dSSRs show a spreading signal (Supplemental Fig. S3A, middle panel). For the longest dSSRs, on occasions we observe a major peak at one edge and a minor signal at the other one (Supplemental Fig. S3A, bottom panel). In some cases, when this asymmetry in H3K4me3 is detected, it prompts the direction of the most expressed strand according to the mRNA detection. This bimodal distribution of H3K4me3 in long dSSRs is consistent with the bidirectional transcription initiation characteristic of *T. cruzi*.

As described above, the extension of the dSSRs in the Dm28c genome is not uniform. To analyze the occurrence of the previously observed patterns in the whole genome, we represented the number of dSSRs for a given length into a histogram and we made an arbitrary division according to most prominent extensions observed (Supplemental Fig., S3B). Therefore, we split the dSSRs into four groups: shorter than 750bp, 750-2500bp, 2500-5000bp, longer than 5000bp. We represented H3K4me3 signal relative to the dSSRs boundaries for these 4 subsets (Fig. 2A). We observed that the trend of H3K4me3 inferred from the observation of individual DGCs is conserved for the whole genome. The group of dSSRs in the range of 0-750bp display a well-defined peak, while for longer regions (750-2500bp, 2500-5000bp) the signal is broader or disperse, suggesting that H3K4me3 is not restricted to a specific location. Instead, alternative positions could be adopted. For those in the longest range (longer than 5000bp) the hint of a second peak is observed, suggesting that among all the alternative positions, those closer to the edges are the most favorable ones. Overall, the longer the region, the greater the number of alternative locations at which H3K4me3 could be found, although some may be more propitious.

**Figure 2.**
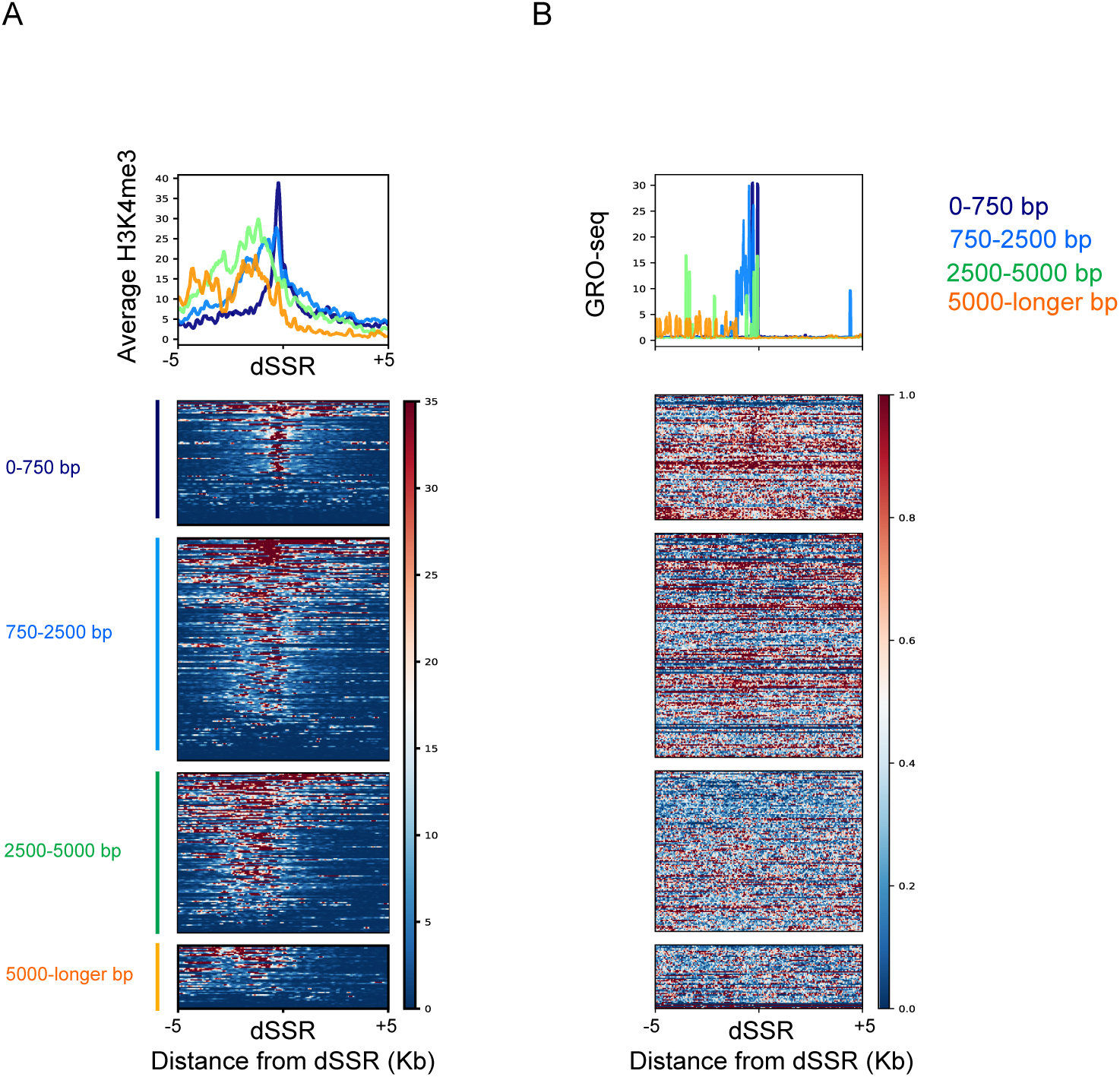
Prevalence of H3K4me3 at highly transcribed DGCs and differential layouts. Average representation and heatmaps for **A)** H3K4me3 and **B)** GRO-seq (SRR28032653) for the four groups of dSSRs defined above: 0-750bp, 750-2500bp, 2500-5000bp, longer than 5000bp relative to the dSSR in a 5Kb window in epimastigotes. Red: high signal; blue: low signal. In both heatmaps, genes are sorted according to the H3K4me3 signal intensities in a descending order.

Then, we wonder whether the different arrangements of H3K4me3 and the associated extension of the dSSRs could be related to the transcriptional outcome. To test this, we analyzed publicly available data sets of GRO-seq (Carvalho de Lima et al., 2024) and we found that, in general, there is an anticorrelation between the signal intensity of the nascent RNA with the length of the dSSRs (Supplemental Fig. S3C). Moreover, we represented GRO-seq signals for the four subsets (Fig. 2B). While heatmap analysis revealed internal variations within each of the four subsets, the average GRO-seq signal highlighted a clear length-dependent trend. Specifically, the shorter dSSR groups (0–750bp and 750–2500bp) displayed robust transcription, while the longest cohorts (>2500 bp) showed reduced average expression.

This result suggests that underneath chromatin organization and epigenetics, there are structural features in the *T. cruzi* genome that might have evolved to guarantee an additional layer of control.

### Distinctive distribution profiles of H3K4me3 within the core and disruptive compartments of the genome

It was previously shown that in *T. cruzi* the core and disruptive genome compartments are associated with different chromatin organization, nucleosome occupancy and gene expression in epimastigotes (Carvalho de Lima et al., 2024; Díaz-Viraqué et al., 2023; Zambrano Siri et al., 2026). To investigate whether DGCs associated with different intensities or patterns of H3K4me3 belong to a specific compartment of the genome, we represented the average H3K4me3 signal relative to the boundaries of the dSSRs for genes located either at the core or the disruptive compartment (Fig. 3A). From a total 460 dSSRs boundaries (coming from 230 dSSRs), 191 are within the core compartment and 269 within the disruptive compartment. A full list for the SSRs located at each compartment could be found at Supplemental Table S3. Our results uphold that H3K4me3 is particularly enriched at dSSRs within the core compartment of the genome (Supplemental Fig. S4A). Representation relative to the dSSRs shows an average enrichment right upstream of the boundaries, while for those in the disruptive compartment the average signal is less intense and more dispersed in the represented window (Fig. 3A). Additionally, H3K4me3 is normally present as a defined peak in the core compartment, while it is broader in the disruptive compartment. This observation is consistent with the wider extension of the dSSRs within this genomic compartment (Supplemental Fig. S4B).

**Figure 3.**
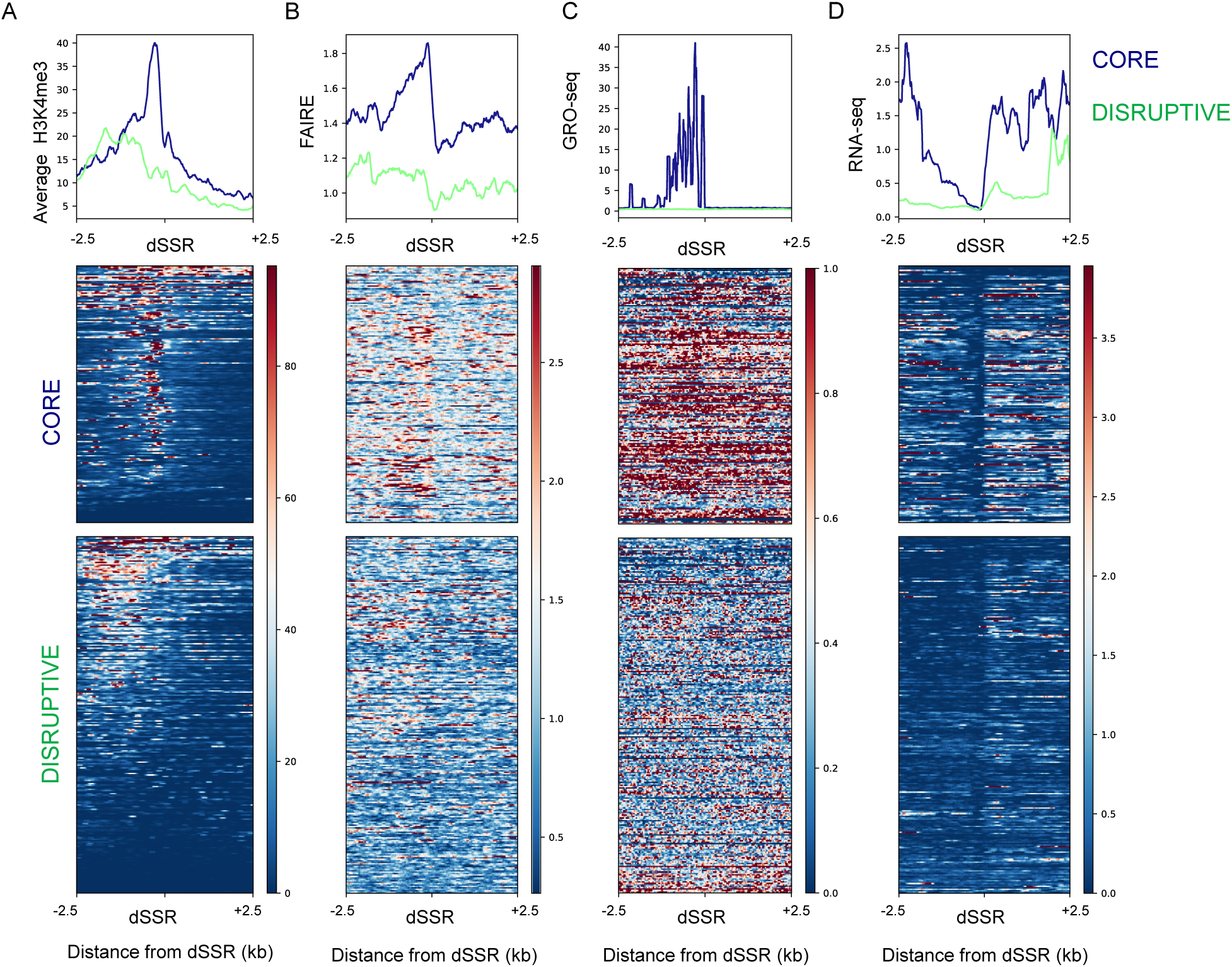
H3K4me3 is more noticeable at dSSRs within the core compartment. Average density (top panels) and heatmaps (bottom panels) relative to the dSSRs in a 2.5 Kb window for DGCs belonging to either the core (blue) or the disruptive (green) compartments of the genome for **A)** H3K4me3 and **B)** FAIRE-seq signals (SRR15902297), **C)** RNA-seq signal (E1), and **D)** GRO-seq (SRR28032653) for a representative replicate of exponentially growing epimastigotes. In the heatmaps genes are sorted according to H3K4me3 signal in a descending order. The same sorting is kept for RNA-seq and FAIRE-seq signals.

To contrast H3K4me3 presence and the above-described patterns with chromatin accessibility for these two subsets of genes, we analyzed public FAIRE-seq data and represented average signals relative to the dSSRs (Fig. 3B). As observed for model organisms, there is a correlation between average H3K4me3 and chromatin accessibility, demonstrating that in *T. cruzi* this epigenetic mark is also associated with open chromatin. Moreover, this correlation is not just an average. Heatmap representations unveil that for most dSSRs located within the core compartment, the most accessible point is right upstream of the edges, coincident with a defined peak of H3K4me3 (Fig. 3A and B, lower panels). On the other hand, when looking at the disruptive compartment, H3K4me3 signal is more dispersed and chromatin is less accessible.

To understand if the lower intensity of H3K4me3 at the disruptive compartment could be explained by lower levels of gene expression, we contrasted the mRNA levels detected by RNA-seq experiments performed with parasites grown in the same experimental conditions. We confirmed that the levels of mRNA are higher for genes within the core-compartment and the presence of H3K4me3 is located at non-transcribed regions but contiguous to actively expressed gene clusters (Fig. 3C). Moreover, to check if H3K4me3 is connected to the transcription rate, we represented GRO-seq available data relative to the dSSRs in both compartments and we found a robust correlation (Fig. 3D). This result highlights an association between H3K4me3, chromatin accessibility and higher expression levels for genes located within the core compartment.

Even for the highly divergent trypanosomes, H3K4me3 appears associated with active transcription, suggesting a conserved role in transcription across different eukaryotic branches.

## Discussion

H3K4me3 is the most ubiquitously conserved epigenetic mark. Although it represents a hallmark of actively transcribed chromatin in eukaryotes, whether it is necessary to initiate active transcription or a result of this, is still a topic of active discussion (Howe et al., 2017). In budding yeast, SET1 methyltransferase is responsible for H3K4 methylation, although this mark is typically repressive. *set1Δ* strains are viable, but they are defective in silencing mating-type loci and telomere regions. They also present growth and sporulation defects (Nislow et al., 1997). It was also described that Pol II guides SET1 as it passes by and leaves this histone mark as a result of its activity (Henikoff and Shilatifard, 2011). In metazoans it was proposed that H3K4me3 has a role in transcriptional pause-release of Pol II (Wang et al., 2023). All this evidence points in the direction of a clear implication in active transcription and genome-wide studies point more evidence in the direction of incorporation upon transcription with a role in directing downstream processes. However, some differences appear among eukaryotes and demand a case-by-case study.

Trypanosomes are highly divergent eukaryotes where gene expression is mainly regulated post-transcriptionally. However, some epigenetic rules seem to be universal, such is the case of some histone PTMs and the presence of the histone variant H2A.Z at the origins of transcription (Maree et al., 2022; McDonald et al., 2022). In particular, H3K4me3 was found to be associated to the transcription start regions, normally at dSSRs, as seen in *T. brucei*, some species of *Leishmania* and *T. cruzi* (Respuela et al., 2008; Siegel et al., 2009; Thomas et al., 2009; Wright et al., 2010). In *L. tarentolae* it was also detected at centromeric regions (McDonald et al., 2022). Similarly, in *L. infantum* H3K4me3 was found at the beginning of DGCs transcribed by Pol II, upstream of RNA genes transcribed by RNA Pol I (rRNA), RNA Pol III (snoRNA, tRNA) and at centromeres (Diotallevi et al., 2025).

In the case of *T. brucei*, it has been described that the starting point and directionality of transcription by RNA Pol II is dependent on a complex called SPARC (Staneva et al., 2022).The positions of such complex in the chromosomes seems to correlate with the positions adopted by the H3K4me3 epigenetic mark here described in *T. cruzi*. Moreover, CRD1, a core component of the SPARC complex, is a chromodomain-containing protein. If CRD1 recognizes H3K4 methylations, then the H3K4me3 could be positioning the SPARC complex and thus defining the TSRs.

Studying epigenetic marks in trypanosomes implies overcoming the fact that their histone sequences are divergent from those of model organisms and in most cases commercial antibodies do not target them. Adding to this complexity, some strains are hard to grow and maintain, making the obtention of the starting material a difficult task. Here, we adapted for the first time the working steps to perform CUT&RUN in epimastigotes and trypomastigotes of *T. cruzi*. This technique requires a small amount of antibody, less starting material and the outcome has less background compared to other widely used techniques, such as ChIP-seq (Skene and Henikoff, 2017).

In *T. cruzi,* H3K4me3 was mapped before the emergence of next generation sequence technologies, through a targeted locus specific experiment, and was shown that in epimastigotes of *T. cruzi* H3K4me3 mapped at dSSRs (Respuela et al., 2008). Here, we refined the analysis by applying CUT&RUN and extended the study to trypomastigotes. We corroborated that H3K4me3 was mainly enriched at dSSRs in both life stages of the parasite (Fig. 1). Moreover, we unveiled that additional H3K4me3 peaks could be found outside dSSRs. Such is the case for minor peaks detected at cSSRs, inside divergent gene clusters (DGCs) or at telomeric regions. A few peaks could also span more than one SSR, or more than one DGC (Supplemental Fig. 1). The role of those peaks with an unusual distribution will require further investigation.

Additionally, comparing its distribution between life stages, we found that H3K4me3 is conserved between epimastigotes and trypomastigotes (Supplemental Fig. 2C). This observation is consistent with our previous knowledge that gene expression regulation along the life cycle occurs mainly post-transcriptionally (Smircich et al., 2015). This result resembles previous observations made in *L. infantum* where H3K4me3 enriched regions are constant between promastigotes and amastigotes (Diotallevi et al., 2025).

From heatmap representations of H3K4me3 we proved a paramount location of this mark at dSSRs in epimastigotes and trypomastigotes, but the signal intensity is not the same at every dSSR along the genome as observed in the heatmaps (Fig. 1A and Supplemental Figure S2 A). This observation raised the question about whether this mark could be related to the levels of expression of the neighboring DGCs. To answer this, we performed RNA-seq in epimastigotes and trypomastigotes and we could observe mRNA signal in the vicinity of H3K4me3 but, in most cases the histone mark does not overlap the transcribed region (Fig. 2A and Supplemental Fig. S3A). This observation suggests a regulatory role for H3K4me3, either as a stimulation for a more efficient transcription or as footprint upon Pol II passage. Further exploration will be required for a more precise mechanistic description.

Furthermore, we inspected individual loci and made average representations revealing that H3K4me3 signal varies with the length of the dSSRs with a negative trend for transcription rate as the length of dSSRs increases (Supplemental Fig. S3C). The possible patterns adopted by H3K4me3 transition from: defined peaks, for short regions; disperse distribution, for intermediate ones; and two peaks near the edges, for the longest- (Fig. 2A and Supplemental Fig. S3A). Moreover, it is worth pointing out that, on occasions, H3K4me3 is mildly boosted on one of the edges of the longest dSSRs, coincidental with a stronger mRNA detection in the associated direction of transcription (Supplemental Fig. S3A, bottom panel). This asymmetry in signaling and the stranded transcription outcome are in agreement with the higher transcription rate reported previously for the positive strand of DNA (Respuela et al., 2008).

This structural organization of the dSSRs is also connected to a previous physical and functional partition described for the genome into core and disruptive compartments (Berná et al., 2018). Analyzing these two subsets we observed that H3K4me3 is labeling dSSRs in both genomic compartments, but the layout is clearly different between them. While the dSSRs located within the core compartment have H3K4me3 confined to the edges, and most of them have a prominent signal, those located within the disruptive compartment present a more diffuse pattern (Fig. 3A). Notoriously, most of the longest dSSR belong to the disruptive compartment while the shortest ones belong to the core compartment (Supplemental Fig. S4B). These observations are also in agreement with differential chromatin accessibility, levels of gene expression and transcription rate detected for the DGCs at core and disruptive compartments (Fig. 3B-D). Additional studies will be required to clarify if this structural organization of the genome is imperative on other layers of regulation.

Overall, our results are consistent with the fact that in trypanosomatids there are no clear promoter sequences. Hence, we suggest that Pol II scans the dSSRs with chances to start transcribing at alternative points along the region, where a longer sequence provide more alternative possibilities to initiate this process. This could also reflect the fact that in a longer sequences there are several fallback positions that a nucleosome can adopt, and in bulk experiments we observe the average patterns of all possible H3K4me3 (Fig. 4). Whether this mark could serve as a buster to make easier successive rounds of transcription or to help in the transition from Pol II initiation to elongation remains unexplored. Single cell experiments together with mechanistic studies should be performed to address this question.

**Figure 4.**
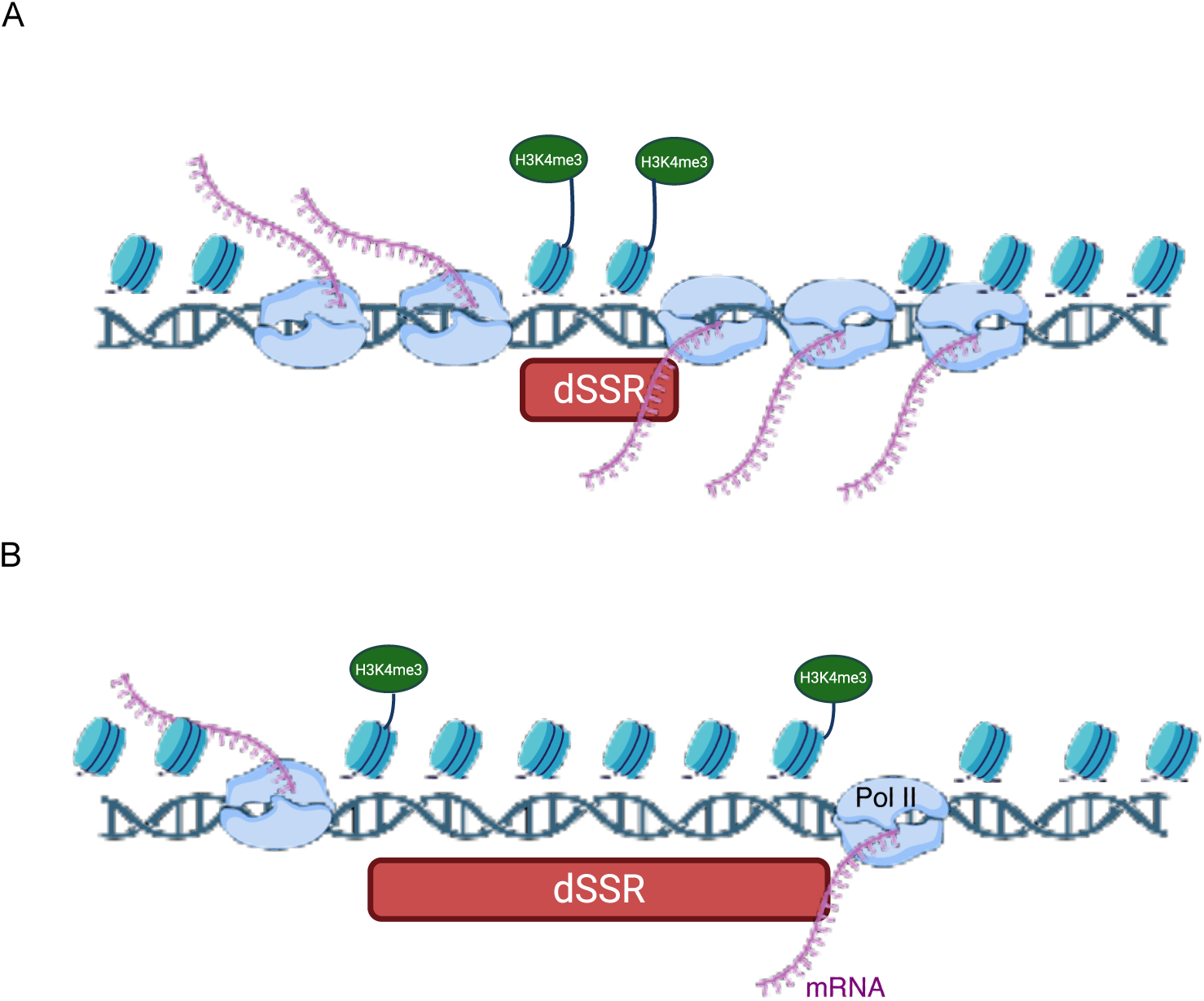
Differential enrichment of H3K4me3 correlates with the genomic span of transcription initiation regions and with transcriptional activity. Schematic representation illustrating the density of H3K4me3 and the transcriptional activity observed at **A)** short and **B)** long dSSRs. Pol II is either more efficiently recruited or more processive at DGCs associated to short dSSRs resulting in more active transcription and higher levels of mRNAs.

## Methods

### Parasites grow and differentiation

Epimastigotes of Dm28c strain were grown at 28°C in LIT medium [5 g/l liver infusion, 5 g/l Bacto-tryptose, 68 mM NaCl, 5.3 mM KCl, 22 mM Na_2_PO_4_, 0.2% (w/v) glucose, 0.002% (w/v) hemin] containing 10% v/v fetal bovine serum (FBS) (NATOCOR, Argentina), 100 units/ml penicillin and 100 μg/l streptomycin. Cell density was maintained between 1×10^6^ and 1×10^8^ cells/ml. Cells were counted using a Neubauer chamber. *Cercopithecus aethiops* (green monkey) Vero cells (ATCC CCL-81) were cultured at 37°C and 5% CO_2_ in Minimum Eagle Medium (MEM, Gibco) supplemented with 5 % FBS, 2 mM Glutamine (Sigma), 100 units/ml penicillin and 100 μg/l streptomycin. To obtain trypomastigotes cells were infected (MOI 10:1 ratio) and 24 h after infection they were washed with Phosphate Buffered Saline (PBS: NaCl 137 mM, KCl 2.7 mM, Na_2_HPO_4_ 10 mM, and KH_2_PO_4_ 1.8 mM) and maintained in MEM. Trypomastigotes present in the supernatant of the culture were harvested by centrifugation at 5000 rpm, for 5 minutes at 72 hours post infection and fixed before storage.

### Cell fixation

As explained above, parasites were collected when reaching 1×10^7^ parasites/ml. Each pellet of cells was resuspended in 1 ml BD FACS Lysis for 15 minutes at room temperature. After that, cells were centrifuged for 5 minutes at 5000 rpm, the supernatant was removed, and pellet was washed with 1 ml of PBS and centrifuged for 5 minutes at 5000 rpm. For storage at −80°C, the pellet was resuspended in 500 ul of DMSO+10% FBS.

### CUT&RUN

We adapted the previously described CUT&RUN protocol to samples of exponentially growing epimastigotes and trypomastigotes of *T. cruzi* (Skene and Henikoff, 2017). A total of 1×10^7^ parasites/ml per sample were subjected to this protocol. Briefly, 10 µl of concanavalin A beads (BioMag®Plus) per sample were washed and resuspended in Binding buffer (HEPES 1 M pH 7.9; KCl 1 M; CaCl_2_ 1 M; MnCl_2_ 1 M). Parasites were washed in PBS1x and resuspended in 900 µl of Wash buffer (HEPES 1 M pH 7.5; NaCl 5 M; Spermidine 2 M; miliQ water; BSA 0.2% and 1x Protease inhibitors MiniO Roche) and 10µl of the previously prepared beads were added to each sample. Then, 50 µl of a 1:50 dilution of anti-H3K4me3 (ab8580) antibody was added. A non-specific anti-IgG antibody (Invitrogen; 02-6102) was used as specificity control. After 1 hour at room temperature with agitation, the samples were washed with Dig Wash Buffer (Wash buffer+ 0.01% digitonin). Subsequently, the samples were resuspended in Dig Wash Buffer and incubated for 10 minutes with agitation at room temperature. Upon removing the supernatant, the beads were resuspended in 50 µl of pA-MNase (CurieCoreTech - Protéines Recombinantes) in Dig Wash buffer and incubated for 10 minutes at room temperature. The beads were resuspended in 50 µl of Dig Buffer and placed on ice. Then,1 µl of CaCl_2_ 10 mM was added to each aliquot, mixed and placed on ice for 30 minutes, shaking lightly every 10 minutes, to activate the pA-MNase. The reaction was stopped by adding 50 µl of Stop buffer (NaCl 5 M; EDTA 0.5 M pH8; EGTA 0.5 M; Digitonin 5%; RNasa A 10 ug/ul), mixed and incubated at 37°C for 10 minutes. Then, the tubes were centrifuged at 15000 rpm at 4°C for 5 minutes and placed in the magnetic rack. The supernatant was transferred to a new tube. 1% SDS was added and incubated for 1 hour at 65°C. After that, 0,5 U of Proteinase K (Thermo Fisher Scientific; EO0491) was added and incubated for 1 hour at 56°C. DNA was purified with QIAquick PCR purification kit (QIAGEN). Sequencing was done with libraries kit for Illumina (NEB#E7645 s/l) and sequenced by Illumina.

### RNA-seq experiments

A total of 1-2×10^7^ parasites of either the epimastigotes or trypomastigotes stage were used. Epimastigotes were harvested during the exponential growth phase, and trypomastigotes were obtained from the supernatant of infected cultures (MOI 10:1 ratio) once sufficient yield and >90% purity was reached, were harvested by centrifugation for 10 minutes at 3000 rpm and washed 3 times with PBS. Cells were collected in 1ml trizol and kept at −80 C until used. RNA was extracted using standard procedures and the quality of the preparation was checked with the Bioanalyzed Qsep1. Strand specific library preparation and sequencing was performed by Macrogen Korea using Illumina TruSeq stranded mRNA library and NovaX.

### Bioinformatic analysis

The sequence quality metrics of raw reads were assessed using FastQC v0.11.9 (https://www.bioinformatics.babraham.ac.uk/projects/fastqc/). During this step, overrepresented adapter sequences were detected and trimmed out using Cutadapt tool v3.5 (https://cutadapt.readthedocs.io/en/stable/) (Martin, 2011) when required. Paired-end reads were aligned using Bowtie2 v2.4.4 (https://bowtiebio.sourceforge.net/bowtie2/index.shtml) (Langmead and Salzberg, 2012) against the recently generated version of Dm28c T2T, generously provided by Dr. Carlos Robello (Greif et al., 2026).

BigWigs files were generated from BAM files as described before (Beati et al., 2023). To build average plots and heatmaps for the disaggregated regions around dSSR or cSSR, computeMatrix and plotHeatmap functions from deepTools version 3.5.1 (https://test-argparse-readoc.readthedocs.io/en/latest/) were used (Ramírez et al., 2014). BED files containing the genomic coordinates for the dSSRs and cSSR were used as regions (detailed in Table S1) and, BigWig files generated for each analyzed dataset were used as score files.

For CUT&Run data, peaks for H3K4me3 were called with MACS2 (Feng et al., 2012) using a threshold for the q-value of 0.05.

## Data access

All raw and processed sequencing data generated in this study have been submitted to the NCBI Gene Expression Omnibus (GEO; https://www.ncbi.nlm.nih.gov/geo/) under accession number GSE334835.

Additionally, a source code is available at: https://github.com/romizambrano/H3K4me3.

## Acknowledgments

We are grateful to Dr. Pablo Smircich for valuable discussions. J.O., S.C.V.L. and G.D.A are members of the Research Career of CONICET. M.d.R.L., A.F.P., R.T.Z.S. and S.C. are Ph.D. fellows and their PhD thesis is carried out at Departamento de Fisiología, Biología Molecular y Celular, Facultad de Ciencias Exactas y Naturales, Universidad de Buenos Aires; I.G. is a Ph.D. fellow and his PhD thesis is carried out at Departamento Química Biológica, Universidad de Buenos Aires.

## References

Beati, P., Massimino Stepñicka, M., Vilchez Larrea, S.C., Smircich, P., Alonso, G.D., Ocampo, J., 2023. Improving genome-wide mapping of nucleosomes in Trypanosome cruzi. PLOS ONE 18, e0293809. 10.1371/journal.pone.0293809

Berná, L., Rodriguez, M., Chiribao, M.L., Parodi-Talice, A., Pita, S., Rijo, G., Alvarez-Valin, F., Robello, C., 2018. Expanding an expanded genome: long-read sequencing of Trypanosoma cruzi. Microb. Genomics 4, 1–41. 10.1099/mgen.0.000177

Carvalho de Lima, P.L., de Sousa Lopes, L., Nunes Rosón, J., Borges, A., Karla Bellini, N., Tahira, A., Santos da Silva, M., Pires, D., Elias, M.C., da Cunha, J.P.C., 2024. “Comprehensive Analysis of Nascent Transcriptome Reveals Diverse Transcriptional Profiles Across the Trypanosoma cruzi Genome Underlining the Regulatory Role of Genome Organization, Chromatin Status, and Cis-Acting Elements.” 10.1101/2024.04.16.589700

Cordon-Obras, C., Gomez-Liñan, C., Torres-Rusillo, S., Vidal-Cobo, I., Lopez-Farfan, D., Barroso-del Jesus, A., Rojas-Barros, D., Carrington, M., Navarro, M., 2022. Identification of sequence-specific promoters driving polycistronic transcription initiation by RNA polymerase II in trypanosomes. Cell Rep. 38. 10.1016/j.celrep.2021.110221

Díaz-Viraqué, F., Chiribao, M.L., Libisch, M.G., Robello, C., 2023. Genome-wide chromatin interaction map for Trypanosoma cruzi. Nat. Microbiol. 8, 2103–2114. 10.1038/s41564-023-01483-y

Diotallevi, A., Amatori, S., Persico, G., Buffi, G., Sordini, E., Giorgio, M., Fanelli, M., Galluzzi, L., 2025. Histone H3 K4 trimethylation occurs mainly at the origins of polycistronic transcription in the genome of Leishmania infantum promastigotes and intracellular amastigotes. BMC Genomics 26, 1–14. 10.1186/s12864-025-11350-1

Feng, J., Liu, T., Qin, B., Zhang, Y., Liu, X.S., 2012. Identifying ChIP-seq enrichment using MACS. Nat. Protoc. 7, 1728–40. 10.1038/nprot.2012.101

Greif, G., Chiribao, M.L., Diaz-Viraque, F., Sanz-Rodriguez, C., Robello, C., 2025. Trypanosoma cruzi Has 32 Chromosomes: A Telomere-to-Telomere Assembly Defines Its Karyotype. 10.1101/2025.03.27.645724

Greif, G., Chiribao, M.L., Díaz-Viraqué, F., Sanz-Rodríguez, C.E., Robello, C., 2026. The complete genome of Trypanosoma cruzi reveals 32 chromosomes and three genomic compartments. BMC Genomics 27, 159. 10.1186/s12864-025-12482-0

Guenther, M.G., Levine, S.S., Boyer, L.A., Jaenisch, R., Young, R.A., 2007. A Chromatin Landmark and Transcription Initiation at Most Promoters in Human Cells. Cell 130, 77–88. 10.1016/j.cell.2007.05.042

Henikoff, S., Shilatifard, A., 2011. Histone modification: cause or cog? Trends Genet. 27, 389–396. 10.1016/j.tig.2011.06.006

Howe, F.S., Fischl, H., Murray, S.C., Mellor, J., 2017. Is H3K4me3 instructive for transcription activation? BioEssays 39, 1–12. 10.1002/bies.201600095

Langmead, B., Salzberg, S.L., 2012. Fast gapped-read alignment with Bowtie 2. Nat. Methods 9, 357–9. 10.1038/nmeth.1923

Maree, J.P., Tvardovskiy, A., Ravnsborg, T., Jensen, O.N., Rudenko, G., Patterton, H., 2022. Trypanosoma brucei histones are heavily modified with combinatorial post-translational modifications and mark Pol II transcription start regions with hyperacetylated H2A. Nucleic Acids Res. 50, 9705–9723. 10.1093/nar/gkac759

Martin, M., 2011. Cutadapt removes adapter sequences from high-throughput sequencing reads. EMBnet.journal 17, 10. 10.14806/ej.17.1.200

Martínez-Calvillo, S., Yan, S., Nguyen, D., Fox, M., Stuart, K., Myler, P.J., 2003. Transcription of Leishmania major Friedlin Chromosome 1 Initiates in Both Directions within a Single Region. Mol. Cell 11, 1291–1299. 10.1016/S1097-2765(03)00143-6

Matthews, K.R., Tschudi, C., Ullu, E., 1994. A common pyrimidine-rich motif governs trans-splicing and polyadenylation of tubulin polycistronic pre-mRNA in trypanosomes. Genes Dev. 8, 491–501. 10.1101/gad.8.4.491

McDonald, J.R., Jensen, B.C., Sur, A., Wong, I.L.K., Beverley, S.M., Myler, P.J., 2022. Localization of Epigenetic Markers in Leishmania Chromatin. Pathogens 11, 1–11. 10.3390/pathogens11080930

Nislow, C., Ray, Evan, Pillus, Lorraine, 1997. SET1, A Yeast Member of theTrithorax Family, Functions in Transcriptional Silencing and Diverse Cellular Processes [WWW Document]. 10.1091/mbc.8.12.2421

Ocampo, J., Carena, S., López, M. del R., Vela, V.S., Zambrano Siri, R.T., Balestra, S.A., Alonso, G.D., 2025. Trypanosomatid histones: the building blocks of the epigenetic code of highly divergent eukaryotes. Biochem. J. 482, 325–340. 10.1042/BCJ20240543

Ramírez, F., Dündar, F., Diehl, S., Grüning, B.A., Manke, T., 2014. deepTools: a flexible platform for exploring deep-sequencing data. Nucleic Acids Res. 42, W187–91. 10.1093/nar/gku365

Respuela, P., Ferella, M., Rada-Iglesias, A., Åslund, L., 2008. Histone acetylation and methylation at sites initiating divergent polycistronic transcription in Trypanosoma cruzi. J. Biol. Chem. 283, 15884–15892. 10.1074/jbc.M802081200

Saha, S., 2020. Histone Modifications and Other Facets of Epigenetic Regulation in Trypanosomatids: Leaving Their Mark. mBio 11, 1–19. 10.1128/mBio.01079-20

Shen, H., Xu, W., Guo, R., Rong, B., Gu, L., Wang, Z., He, C., Zheng, L., Hu, X., Hu, Z., Shao, Z.-M., Yang, P., Wu, F., Shi, Y.G., Shi, Y., Lan, F., 2016. Suppression of Enhancer Overactivation by a RACK7-Histone Demethylase Complex. Cell 165, 331–342. 10.1016/j.cell.2016.02.064

Siegel, T.N., Hekstra, D.R., Kemp, L.E., Figueiredo, L.M., Lowell, J.E., Fenyo, D., Wang, X., Dewell, S., Cross, G.A.M., 2009. Four histone variants mark the boundaries of polycistronic transcription units in Trypanosoma brucei. Genes Dev. 23, 1063–1076. 10.1101/gad.1790409

Skene, P.J., Henikoff, S., 2017. An efficient targeted nuclease strategy for high-resolution mapping of DNA binding sites. eLife 6, 1–35. 10.7554/eLife.21856

Smircich, P., Eastman, G., Bispo, S., Duhagon, M.A., Guerra-Slompo, E.P., Garat, B., Goldenberg, S., Munroe, D.J., Dallagiovanna, B., Holetz, F., Sotelo-Silveira, J.R., 2015. Ribosome profiling reveals translation control as a key mechanism generating differential gene expression in Trypanosoma cruzi. BMC Genomics 16, 443. 10.1186/s12864-015-1563-8

Staneva, D.P., Bresson, S., Auchynnikava, T., Spanos, C., Rappsilber, J., Jeyaprakash, A.A., Tollervey, D., Matthews, K.R., Allshire, R.C., 2022. The SPARC complex defines RNAPII promoters in Trypanosoma brucei. eLife 11, 1–23. 10.7554/eLife.83135

Strahl, B.D., Allis, C.D., 2000. The language of covalent histone modifications. Nature 403, 41–45. 10.1038/47412

Thomas, S., Green, A., Sturm, N.R., Campbell, D.A., Myler, P.J., 2009. Histone acetylations mark origins of polycistronic transcription in Leishmania major. BMC Genomics 10, 1–15. 10.1186/1471-2164-10-152

Tyler, K.M., Engman, D.M., 2001. The life cycle of *Trypanosoma cruzi* revisited. Int. J. Parasitol., The Third Internet Conference on Salivarian Trypanosomes and Trypanosomatids 31, 472–481. 10.1016/S0020-7519(01)00153-9

Wang, H., Fan, Z., Shliaha, P.V., Miele, M., Hendrickson, R.C., Jiang, X., Helin, K., 2023. H3K4me3 regulates RNA polymerase II promoter-proximal pause-release. Nature 615, 339–348. 10.1038/s41586-023-05780-8

Wang, H., Helin, K., 2025. Roles of H3K4 methylation in biology and disease. Trends Cell Biol. 35, 115–128. 10.1016/j.tcb.2024.06.001

Wedel, C., Förstner, K.U., Derr, R., Siegel, T.N., 2017. GT-rich promoters can drive RNA pol II transcription and deposition of H2A.Z in African trypanosomes. EMBO J. 36, 2581–2594. 10.15252/embj.201695323

Wright, J.R., Siegel, T.N., Cross, G.A.M.M., 2010. Histone H3 trimethylated at lysine 4 is enriched at probable transcription start sites in Trypanosoma brucei. Mol. Biochem. Parasitol. 172, 141–144. 10.1016/j.molbiopara.2010.03.013

Zambrano Siri, R.T.Z., Beati, P., Inchausti, L., Smircich, P., Alonso, G.D., Ocampo, J., 2026. Beyond the transcript: Chromatin implications in trans-splicing in Trypanosomatids. PLOS ONE 21, e0343367. 10.1371/journal.pone.0343367

