## Supplemental Figures for "H3K4me3 exhibits length-dependent deposition patterns at transcription initiation regions in *Trypanosoma cruzi* and correlates with transcriptional activity"

**Supplemental Figure 1**

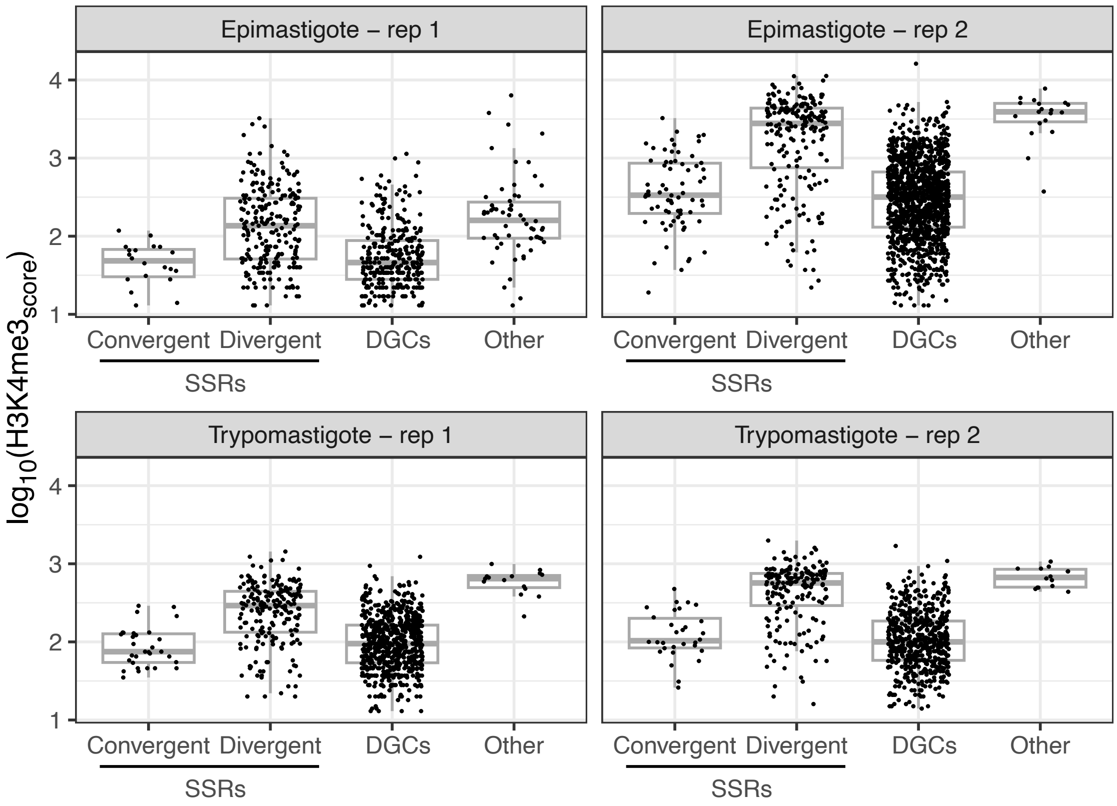


**S1 Figure. H3K4me3 peaks distribution throughout the genome**

Score of H3K4me3 peaks detected by MACS2 at different genomic locations. Convergent or divergent strand switch regions (SSRs), divergent gene clusters (DGCs) or Other are represented. Others include peaks detected at: telomeric regions, that spans more than one SSR, or more than one DGC. Y axis represents the intensity of the peaks for each sample tested.

#### Supplemental Figure 2

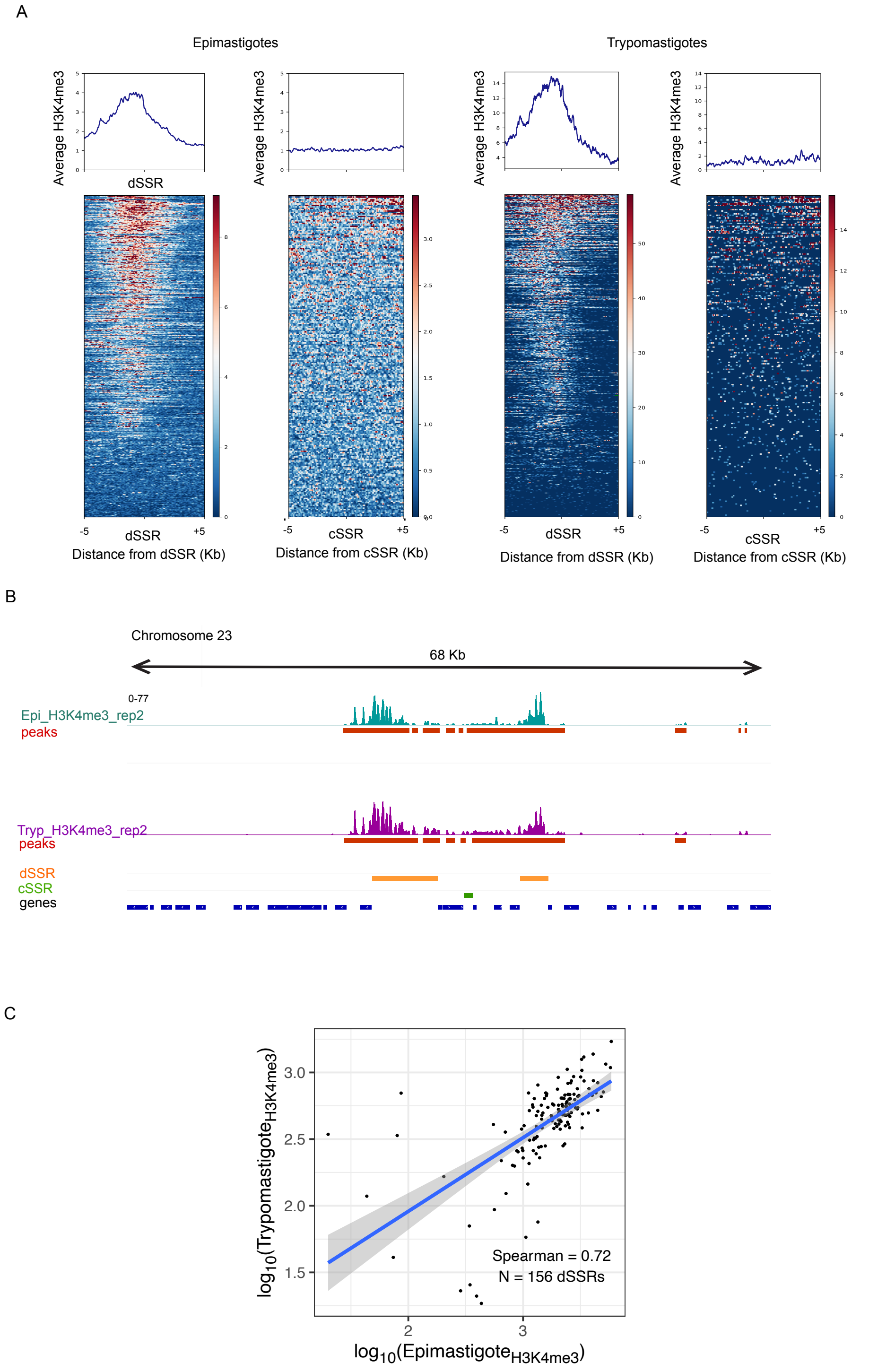

**S2 Figure. H3K4me3 is a signature of dSSRs at epimastigotes and trypomastigotes**

**A)** Average H3K4me3 density (top panels) and heatmaps (bottom panels) relative to the dSSR (left) or cSSR (right) in a 5 Kb window in epimastigotes and trypomastigotes for a replicated experiment. In the heatmaps genes are sorted according to H3K4me3 signal in a descending order. For those represented relative to cSSR, annotations for DGCs shorter than 5 Kb were removed to avoid the misleading observation of the neighbor dSSR. Red: high signal; blue: low signal, green: missing data. **B)** IGV image showing a region of chromosome 23 of T2T Dm28c genome displaying H3K4me3 tracks for a representative replicate experiment in epimastigotes (cyan) and trypomastigotes (purple) with their respective narrow peaks (red) generated with MACS2. The annotated coordinates for dSSRs (orange) and cSSRs (green) for the used genome are included as a reference. An example of these regions that are in proximity from each other is illustrated. **C)** Scatterplot of H3K4me3 signal in epimastigotes (x-axis) versus trypomastigotes (y-axis). The H3K4me3 signal was quantified for each dSSR by summing the scores of MACS2-identified peaks that fully or partially overlap the region. Signals were averaged across biological replicates for each life stage. dSSRs lacking detectable peaks in one or both life stages were excluded from the analysis.

### Supplemental Figure 3

A

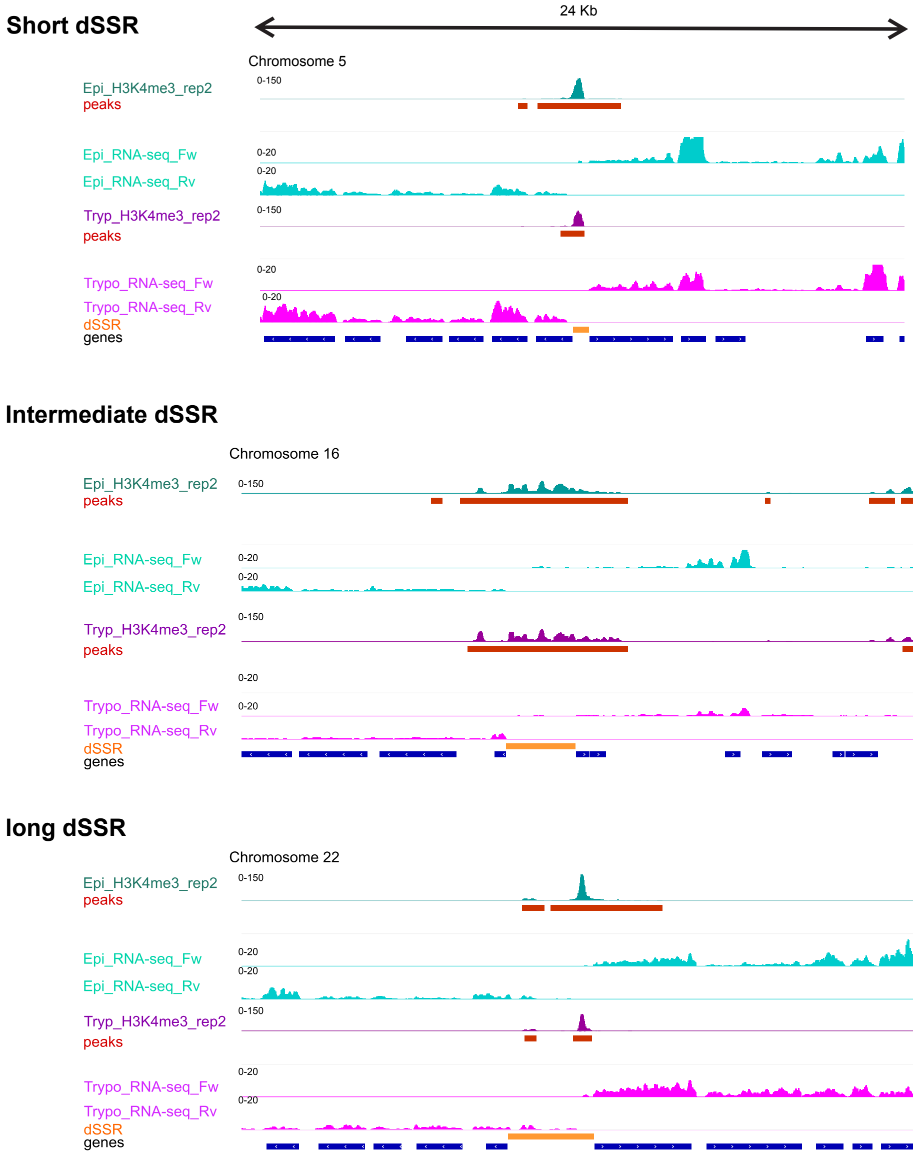


B

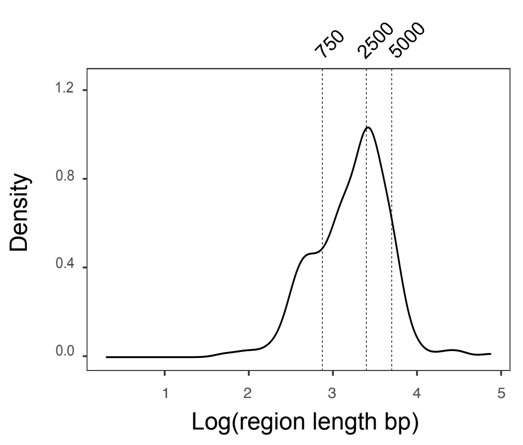


C

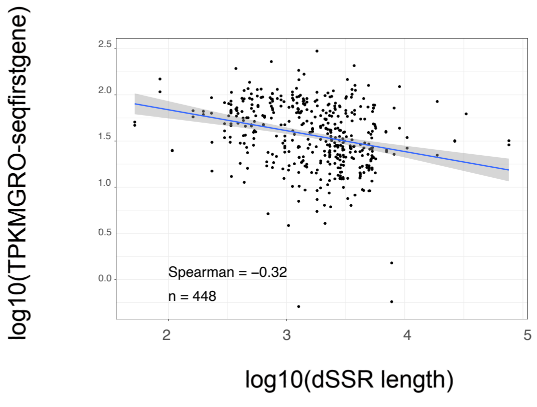

**S3 Figure. H3K4me3 display different patterns related to the extension of the dSSR**

**A)** IGV image showing examples of dSSR, that belongs to the 0-750 bp (top), 2500-5000bp group (middle), and longer to 5000bp (bottom panel), in a fix window of 24kb of the T2T Dm28c genome displaying H3K4me3 and RNA-seq signals for one representative replicate of epimastigotes (cyan) and trypomastigotes (purple) with their respective narrow peaks (red) generated with MACS2. Annotated dSSR coordinates for the used genome are included for reference (orange). **B**) Density plot showing the distribution of dSSRs lengths. **C)** Scatterplot representing the TPM for GRO-seq signal of the first gene of the DGCs in epimastigotes (y-axis) versus the length of the dSSRs (x-axis) in log10 scale.

**Supplemental Figure 4**

A

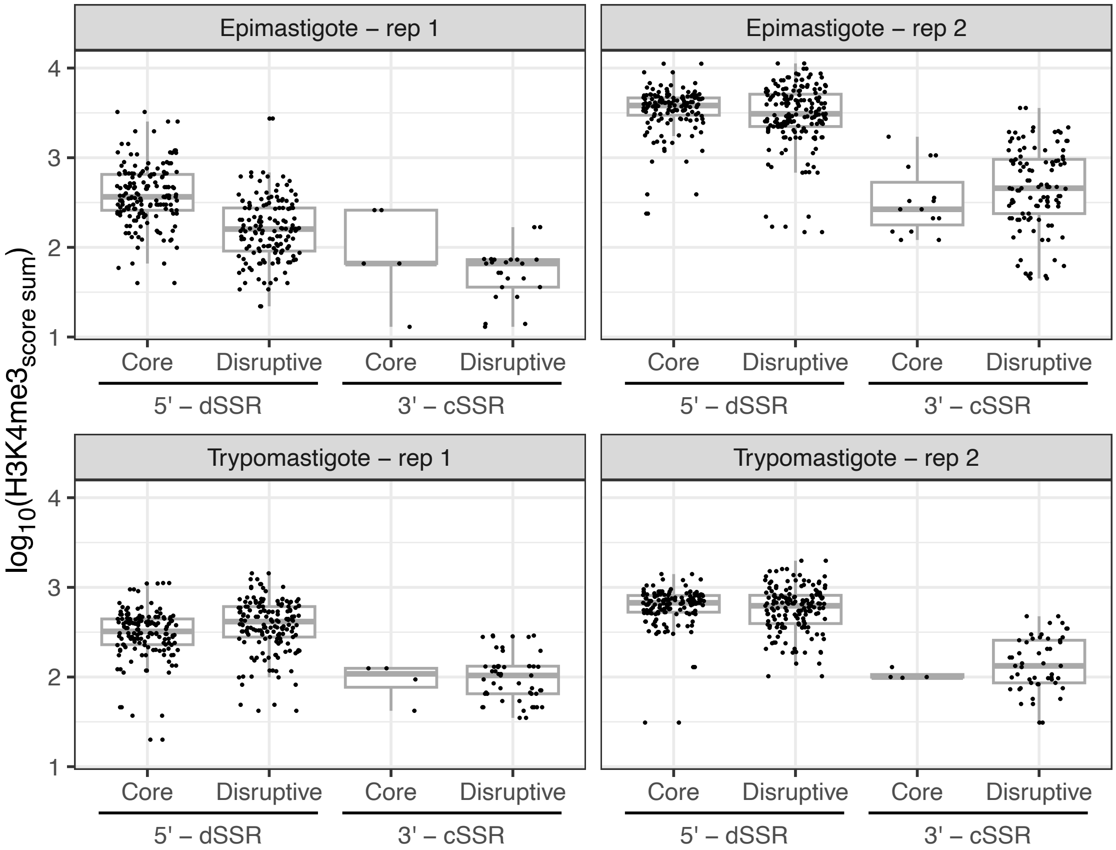


B

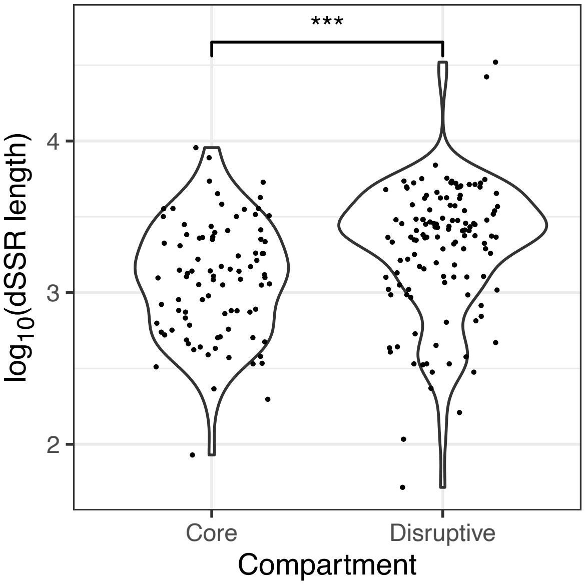

**S4 Figure. H3K4me3 distribution at core and disruptive compartments of the genome**

**A)** Intensity of H3K4me3 peaks detected by MACS2 at different genomic locations. The boundaries of convergent (cSSRs), or divergent strand switch regions (dSSRs), that fall into either the core or the disruptive compartment for each sample tested. **B)** The y axis shows the dSSRs length (in log10 scale) for those falling in the core or disruptive compartment. Those dSSRs falling in both compartments were excluded from the plot. The significance level corresponds to a p-value < 0.001 in a Student's t-test.
